# Predicting recoverability of collapsed food webs through perturbation and dimension reduction

**DOI:** 10.1101/2024.07.09.602684

**Authors:** Swastik Patnaik, Gaurav Baruah

## Abstract

Biodiversity collapse, driven by escalating environmental changes, poses significant threats to ecosystem stability and the provision of essential ecosystem services. Understanding the recoverability of collapsed food webs thus is crucial for devising effective conservation strategies. This study delves into the theoretical exploration of the recoverability of food webs from a collapsed state. Through simple tools like dimension reduction, propagation of species-specific perturbation, and dynamical simulations, we explore whether simple tri-trophic food webs can be recovered from a collapsed state. Our study examines in detail the topological features of predator-prey food webs that could either facilitate or impede their recovery. We demonstrate that the recoverability of complex food webs can be predicted by using a simple dimension-reduced model, with certain structural factors that could constrain the full recovery of collapsed food webs. Furthermore, dynamic simulations also highlighted the significance of topological features such as connectance and the number of predator links in determining recoverability. Our dimension-reduced modeling framework offers insights into the feasibility of restoring entire complex predator-prey networks through species-specific interventions. This study contributes to a deeper understanding of ecosystem resilience and aids in the development of targeted conservation strategies.

## 1 Introduction

With an increase in loss of biodiversity at an alarming rate, conserving species diversity has thus become increasingly important. Complex ecological networks are increasingly at the risk of catastrophic transitions which often are a result of increasing anthropogenic pressures (Borrvall & Ebenman 2006, Lindegren *et al*. 2012, Dunne & Williams 2009, Kratina *et al*. 2012, Lenton *et al*. 2008, Hutchings & Reynolds 2004, Scheffer *et al*. 2001, Cardillo *et al*. 2005). As the threat of species loss increases, there is also a concurrent risk of loss of important ecosystem services and functions associated with it (Smith & Schindler 2009, Scheffer *et al*. 2001). Some examples of transitions of ecological networks are the collapse of fisheries (Dakos *et al*. 2019, Pinsky *et al*. 2011, Hutchings & Reynolds 2004), or the collapse of vegetation leading to a desert state (Kéfi *et al*. 2007), or the collapse of lake ecosystem from a clear state to a turbid state (Scheffer *et al*. 2001). Transitions such as these are a result of changes in populations of species which are a direct or indirect consequence of changes in the external environment (Baruah & Lakämper 2024). As changes in populations in such complex networks occur due to an external environmental perturbation, it can ripple through an ecosystem and cause undesirable changes (Gilarranz *et al*. 2017). Because such complex ecological networks are interconnected, the extinction of a species can have a cascading impact on the resilience and structure of ecosystems (Hens *et al*. 2019). Thus, forecasting future changes in species composition in complex food web communities due to anthropogenic pressures remains a challenge (Lever *et al*. 2020, Medeiros *et al*. 2023).

Forecasting changes in ecological communities over time has been a challenge but an important goal for biodiversity conservation (Dakos *et al*. 2012, Scheffer *et al*. 2009, Baruah *et al*. 2019). A challenging aspect of this is that it requires tracking complex ecological interactions over time in addition to monitoring species dynamics embedded in these communities (Baruah *et al*. 2022, Deyle *et al*. 2016). In addition, these interactions can change due to shifts in population densities of species as a result of shifts in the external environment. As the external environment keeps changing, complex communities could shift to undesirable states (Baruah 2022, Lever *et al*. 2014, Yacine *et al*. 2021). Promising tools that have been developed to forecast such changes include phenomenological tools such as early warning signals and empirical dynamical modelling (Deyle *et al*. 2016, Dakos *et al*. 2012, Baruah *et al*. 2019, Sugihara *et al*. 2011, Arkilanian *et al*. 2020). Both of these tools require the monitoring of populations embedded in complex communities extensively. In case species intertwined in these complex communities do collapse, it could trigger a cascade in such communities (Dunne & Williams 2009, Dobson *et al*. 2009). For instance, the loss of predators could trigger extinction cascades in a complex food web community. Theoretical as well as empirical studies, have suggested that loss of species could lead to secondary extinctions (Ebenman *et al*. 2004, Eklöf *et al*. 2012, Fowler & Ruokolainen 2013, Borrvall & Ebenman 2006, Donohue *et al*. 2017, Terborgh *et al*. 2001, Springer *et al*. 2003). Once such complex co-extinctions lead to a collapse of whole food webs (Gilljam *et al*. 2015), it could be difficult to recover such communities. This is because more often we might not have information about such complex communities at the brink of collapse (Baruah & Wittmann 2023). In addition, due to negative feedback among species at different trophic levels, restoring a species might not always lead to a successful recovery. Thus, recovering an entire ecological network from an undesirable collapse state is a problem that has been overlooked. Whether a complex ecological community could even be recovered from a collapsed state needs to be understood (Baruah & Wittmann 2023).

The high-dimensional nature of complex dynamical systems poses a challenge to evaluate how the system behaves near the vicinity of a tipping point. A complex ecological system such as a predator-prey food web consists of many interaction species at different trophic levels. Once such a dynamical system collapses, whether recovery of such a system is possible is a challenging problem to address (Dobson *et al*. 2009). One way to understand whether a complex high-dimensional network could be recoverable would be to use the approach of dimension reduction (Jiang *et al*. 2018). Because of many interacting species, the corresponding state-space dimensionality of predator-prey food webs could generally be high, making simple theoretical predictions and insights extremely difficult. Dimension reduction has been used in understanding the behavior of tipping points specifically in mutualistic bipartite ecological networks (Jiang *et al*. 2018). By reducing the dimension of trophic food web networks, one could effectively understand whether a system could potentially recover from a collapsed state. For instance, a complex food web of *n*-species thus has *n* dimensions whose dynamics are complex due to prevalence of non-linearity and interdependence of species on each other in such systems. Thus, if such a system of *n* dimensions could be collapsed into *s* dimensions where *s << n*, then it would be simpler to effectively understand and approximate a complex system’s dynamics, and evaluate the parameters that could potentially impact the stability and resilience of such systems. Thus, dimension reduction entails in reducing the number of possible dimensions to effectively a few dimensions to better understand and predict the dynamics of complex systems(Jiang *et al*. 2018, Gao *et al*. 2016).

The persistence of a species within a intricate food web is influenced by its ability to adapt to external environmental changes and its interactions with other species. However, in complex food webs, the energy flows to different trophic levels are determined by a host of factors including but not limited to biotic interactions such as prey-predation relationship, competition within trophic levels, omnivory etc.(Petchey *et al*. 2008, Dunne *et al*. 2002, Dunne & Williams 2009, Staniczenko *et al*. 2010). All these biotic and abiotic factors would contribute to the resilience of a foodweb to enviromnental perturbation. Once such a foodweb collapses due to environmental perturbation crossing a certain threshold, restoration of such collapsed food web would depend on these factors along with structural properties of the community. We thus propose to investigate the recoverability of collapsed food webs by utilizing the phenomenon of perturbation propagation and dimension reduction of such trophic networks. For instance, previous studies evaluated the impact of topological features of a network and trait variation in species on the revival of collapsed networks using perturbation on a single node of the network (Hens *et al*. 2019, Baruah & Wittmann 2023). Due to positive feed-back loops in mutualistic plant-pollinator networks, a positive perturbation on a single species can in some cases spread over entire networks, leading to a successful revival (Baruah & Wittmann 2023). In contrast, prey-predator food web networks majorly consist of interactions that are negative in nature, such as competition within a trophic level or predation across a trophic level. Due to this, the successful recoverability of an entire trophic network through positive perturbation of a species or groups of species is therefore not guaranteed. However, using dimension-reduction of a complex food web, one could possibly predict whether recoverability of a collapse food web through perturbation could be a possibility. In this study, we take a theoretical perspective in understanding the recoverability of collapsed predator-prey foodwebs (Fig 1). We address whether simple trophic food webs could be recovered from a collapsed state by using dimension reduction, propagation of species perturbation, and dynamical simulations (Fig. 1). We try to understand the topological features of predator-prey food webs that could either hinder or promote the recovery of collapsed food webs. We show that whether a complex food web could be recovered can be predicted through dimension-reduction, and that certain structural factors can constrain the recoverability. We then, using dynamic simulations, show that recoverability of collapsed food webs depends particularly on topological features such as connectance, number of trophic links. Our reduced modeling framework of predator-prey food webs could serve as an indicator of whether entire complex predator-prey networks could be recovered through species-specific intervention.

**Figure 1:**
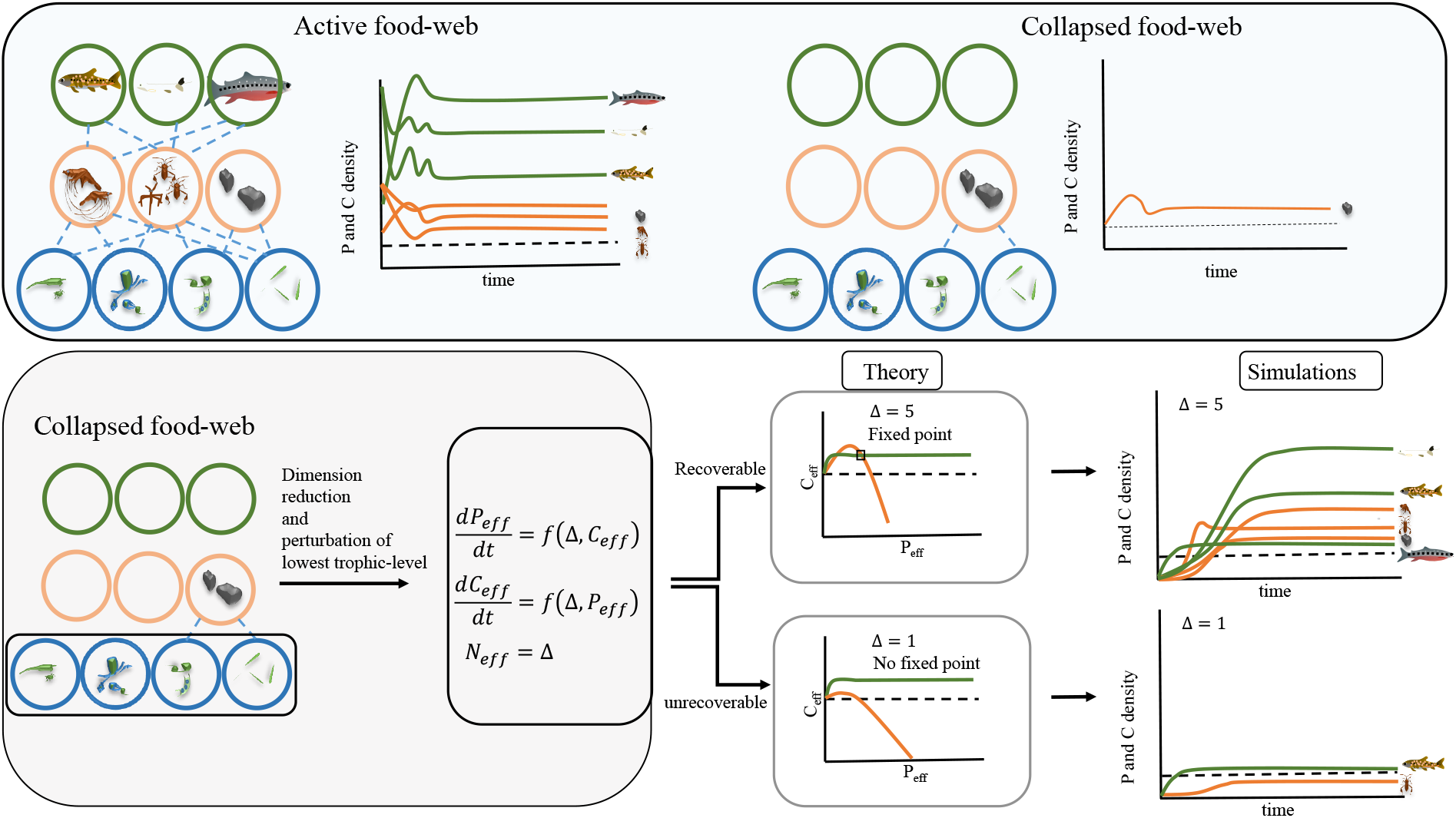
Graphical illustration of the food web collapse and recoverability. A) Illustration of an active high-functioning tri-trophic food web. B) A collapsed food web where only a single primary consumer has a density above a detection extinction threshold (a horizontal dotted line). (C) The same high-dimensional tri-trophic food web could be reduced to a 2-D model based on information from adjacency feeding links of the collapsed food webs. Once the effective 2-D model is derived, it can be used to understand whether perturbing the lowest trophic level could lead to recoverabilty of the entire food web, and this can be easily assessed by analysing the 2-D equation and assessing the emergence of a fixed point based of intersection of the two nullclines of the 2-D equation. Furthermore, this 2-D equation could be further used to see whether a reduced model could accurately match the recovery dynamics of a collapsed food web. D) Then, using simulations of the full model, we assessed whether results of the 2-D reduced model matches those with simulations of a full dynamical food web model.

## 2 Methods and Models

For simplicity we restricted our modelling and analysis to three trophic levels. We simulated theoretical food webs that ranged in connectance values from 0.08, to 0.4. Connectance of a food web is the number of interaction links observed out of number of possible interactions. In these tri-trophic food webs, each species competes with others within a trophic level. The strength of this intraspecific competition was highest in the basal resources, and lowest for consumers and top predators. In other words, primary and secondary consumers had the lower intraspecific competition strength than primary basal producers (details in table 1).

**Table 1:** Parameters and their values with the description used in the study. *U*[*a, b*] is the uniform distribution between *a* and *b*.

| Symbol | Description |
| --- | --- |
| $b_i^{C,P}$ | death rate for primary and secondary consumers; -0.1 for primary consumers, -0.05 for secondary consumers. |
| $a_{ij}^C$ | Interspecific competition was fixed at 0.1 and intraspecific competition of primary consumers was fixed at 0.5; Secondary consumers did not have interspecific competition but only intraspecific competition which was fixed at 0.1 |
| $a^P$ | Intraspecific competition for predator species fixed at 0.05. |
| $\alpha$ | effective competition of the reduced model which was fixed at 0.1 for consumers and secondary consumers. |
| $e$ | conversion efficiency of all consumers and secondary consumers, fixed at 0.5. |
| $\alpha$ | attack rate of secondary and primary consumers which was fixed at 1.25 |
| $h$ | handling time of consumers and secondary consumers which was fixed at 0.25. |
| $A_{ij}$ | adjacency matrix of feeding links between consumers, basal species and secondary consumers or top predators |
| $\Delta$ | Equilibrium density of the lowest trophic level used as a perturbation parameter for recoverability of food webs. |
| $N_i, C_i, P_i$ | Population density of basal resources, primary consumers and secondary consumers respectively. Initial density of all species was fixed at 0.000001. |

### 2.1 Theoretical food webs

The theoretical food webs were generated through either pyramidal or probabilistic niche-based food webs (PNM) see Williams *et al*. (2010), Gilljam *et al*. (2015). In the main-text we show results for the food webs generated through the pyramidal method. For the pyramidal food webs we generate communities ranging from 12 to 24 species distributed over three trophic levels. The ratio of species in the three trophic levels was 5:3:2, and all primary and secondary consumers had at least one feeding link. Connectance of the food web was randomly drawn from a range of 0.08 to 0.4. Mortality or death rates for primary consumers was fixed at -0.1, and -0.05 for secondary consumers resulting in lower rates for consumers at higher trophic levels. This was in relation to the fact that larger body size is related to higher trophic levels and usually confer to lower death rates (Kaneryd *et al*. 2012). Interspecific competition was present in the basal resources and was randomly drawn from a uniform distribution U[0.1,0.5], interspecific competition was fixed at 0.1 for the primary consumers and there was no interspecific competition modeled for top secondary consumers. Intraspecific competition for basal resources was fixed at 1, for primary consumers it was fixed at 0.5, and secondary consumers it was fixed at 0.1. These competition coefficients were used to signify that weak competition at higher trophic levels as indicated in previous studies (Petchey *et al*. 2008, Cohen *et al*. 2003). Omnivory links were not allowed. On the other hand, the PNM method uses a Gaussian niche-based probability that a species feeds on another. The closer the niche position of a resource is to a consumer, the higher the chance of feeding as detailed in Williams *et al*. (2010). Ominvory links that were generated were discarded. Inter and intraspecific competition was similar to as used in building the pyramidal food webs. The numerical PMN and pyramidal food webs generated represent a wide variety of structures in agreement to what is found in real networks (Johnson *et al*. 2014). Please refer to supplementary section 3 for results of two food webs generated through PMN method.

### 2.2 Modeling framework

Dynamics of tri-trophic food webs are given by Rosenzweig-MacArthur model with type-2 functional curves as (Kaneryd *et al*. 2012, Rosenzweig & MacArthur 1963):

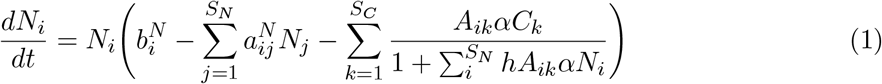

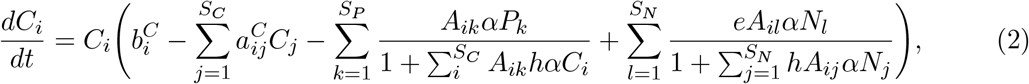

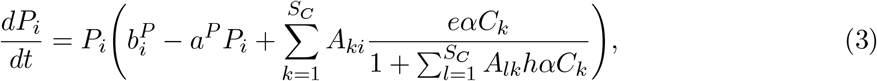

Here, *N*_*i*_, *C*_*i*_, *P*_*i*_ are the density of basal resources, primary consumers, and top predators respectively. 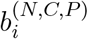 is the per capita growth rate for basal producers, or mortality rate for primary and secondary consumers (see table 1 for details of parameter values); 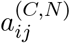 captures the per-capita inter and intraspecific competition within a trophic level. *S*_*N*_ is the group of species being the competitors of basal species, *S*_*C*_ is the group of species that feeds on basal species i.e., the primary consumers, *S*_*P*_ is the group of species that feed on the consumer species (secondary consumers or top predators). **A** is the adjacency matrix of feeding links between a primary basal species, primary consumers, and secondary consumers. Its value is either 0 or 1, where 0 would indicate no feeding link and 1 would indicate an existence of a feeding link. *α* captures the attack rate of primary and secondary consumers, *e* captures the conversion efficiency or the rate at which a resource is efficiently converted to a new consumer, *h* is the handling time. Equation 1,2,3 captures the dynamics of a tri-trophic food web with type 2 functional response. The structure of the food web network is imposed by the adjacency feeding matrix *A*_*ik*_ generated either by PMN or pyramidal method.

### 2.3 Perturbation regime, and dimension reduction

The starting point of our analysis are food webs that have collapsed. We thus define a food web to be collapsed if all primary and secondary consumer species densities have densities *N*_*i*_ *<* 0.00001 irrespecitve of the presence or absence of basal species. The collapse of such a food web could be due to numerous reasons including external environmental change, mortality imposed due to a harvesting regime etc. Our focus was not in evaluating what could have led to the collapse of such food webs. Instead, we wanted to evaluate whether a collapsed food web could be recovered. Thus, we needed to define the recoverability of a collapsed food web. We define recoverability of a collapse food web as the conditions that lead to persistence of at least 70% of the primary or secondary consumer species at densities above 0.1. Ecological systems can transition to undesirable states due to intrinsic or extrinsic factors (Baruah 2022, Scheffer 2009). Restoring original environmental conditions, such as habitat restoration (Kaiser-Bunbury *et al*. 2017, Dobson *et al*. 2009, Downing *et al*. 2012), could lead to recoverability of communities. Here, similar to this, we dictate the control of the lowest trophic level i.e., the basal producer species, and evaluate whether perturbing the lowest trophic level could lead to the recovery of the primary and secondary consumers.

Our analysis is structured as follows. We first assign a perturbation regime to the basal producers and try to analytically through dimension-reduction evaluate whether food webs of different structures could be recovered. Next, using dynamical simulations, we try to assess whether the predictability of recovery dynamics through dimension reduction matches with our simulations, and further evaluate the impact of different food web structure on recoverability. Since we were interested in evaluating whether a food web network could be recovered through perturbation of the lowest trophic level, we first try to reduce complex food webs to two-dimensional state space. Thus a three-trophic food web of n-species with n-dimensions effectively changes to two dimensions capturing the dynamics of effective primary and secondary consumers. In addition, the dynamics of the lowest trophic level of primary producers are controlled through a constant perturbation i.e.,

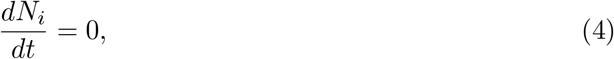

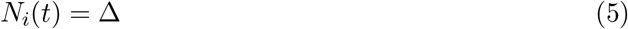

where, all basal producer species *i* are fixed to be at a constant density of Δ. This meant that we control the density of all basal species to be at a density Δ. However, we also apply a continuous rate of forcing, where we positively force all the basal species at a rate *β* and compare this with the dimension-reduction method (see below and see supplementary 1).

Effectively, equation 2 and 3 can be transformed to only primary consumers and secondary consumer equations as:

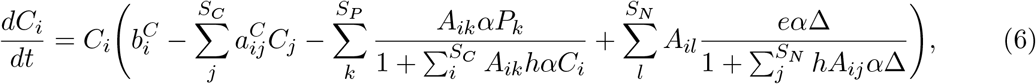

Note Δ in equation 6. However, the secondary consumer equation remains the same as equation 3.

In this context, the dynamics of both primary and secondary consumers are captured by a system of ordinary differential equations. The total number of equations corresponds to the number of primary and secondary consumers present in the system. Analyzing such a high-dimensional state space poses challenges, especially when trying to comprehend the overall resilience of the system. Predicting whether a complex food web that has collapsed could recover under specific perturbation conditions affecting basal producers might be more feasible with knowledge of stability and resilience equations for all consumers and secondary consumers involved. While it is analytically impossible to find out the stable and unstable states of all the primary and secondary consumers given a certain rate of perturbation Δ, we can, however, find out the fixed points of both the primary and secondary consumers by having some kind of aggregate functions dictating the collective dynamics of the primary and secondary consumers through dimension-reduction method (Jiang *et al*. 2018). In the dimension reduction method we aggregate equations 6 and 3 from *n*-state space (where *n* is the number of species) to two state-space model dictating the dynamics of effective primary and effective secondary consumer given the basal species are at a density of Δ. Our goal here was to evaluate whether a simple reduced two-dimensional model could effectively capture the recovery dynamics of a complex tri-trophic food web and thus could inform us about recoverability in general (please see the result section for the dimension reduction model).

### 2.4 Food web topology and species-specific forcing

To explore the relationship between food web structure and the recoverability of collapsed food webs, we generate food webs of different connectance. Specifically, we show in the main text the food webs generated through pyramidal method, and for food webs generated through PMN method, please refer to supplementary appendix 2. We generated pyramidal food webs of different connectance, C, that ranged 0.09 to 0.4 with the total number of species being between 12 and 24. As food webs are generated with different connectance for a particular network size of either 12, 14, 15, or 24, we ensured that each consumer has at least one trophic link. Furthermore, the ratio of species in each of the trophic levels was 5:3:2. We thus evaluated how variation in food web structure could impact the recoverability of collapsed food webs by positive perturbation of the lowest trophic level Δ. When assessing the recoverability of food webs, we ensured that the starting density of all secondary and primary consumer species was below 0.00001. In addition, we also varied the proportion of secondary consumer trophic links to assess the impact of predation on the recoverability of collapsed food webs. The proportion of secondary consumer links varied from as low as 0.2 to 1, where 0.2 would mean out of all possible feeding links of secondary consumer on primary consumer, only 20 % of links are achieved, and 1 would mean out of all possible secondary consumer to primary consumer feeding link, all links are achieved. Thus, we assessed how variation in secondary consumer trophic links and variation in food web connectance impacted food web recovery for a certain Δ.

We assumed that the lowest trophic level i.e., the basal producers were kept at a constant density of Δ while assessing the recovery of secondary and primary consumers. This meant that all basal species were kept at a density of Δ. We however also evaluated how many basal species were needed to be kept at a certain density Δ that leads to the recoverability of food webs. We thus varied Δ from 0 to 5 and varied the number of basal species that was kept at density Δ. This we did for three different levels of connectance, C, from 0.09, 0.15, and 0.25, and for a food web size of 24 species.

## 3 Results

### 3.1 Dimension reduction, perturbation, and recovery dynamics

We can reduce the dimension of equation 6 and 3 to effectively two equations. The idea would be to aggregate information about the entire network by an effective parameter. Effective densities of primary and secondary consumer thus can be represented by *C*_*e*_ and *P*_*e*_. For simplicity we ignore interspecific competition as we parameterised the model with intraspecific competition being significantly larger than interspecific competition. Hence, the terms capturing interspecific competiton in equation 6 and 3 can effectively be reduced to

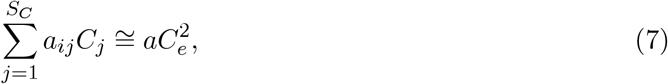

and

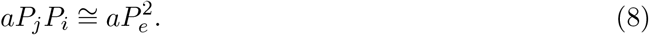

where, *P*_*e*_ is the effective density of top predators, and *C*_*e*_ is the effective density of primary consumers. Using the degree-weighted method following(Jiang *et al*. 2018), the predation of primary consumer on the basal species (third term of equation 4), can be written as

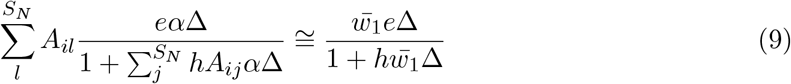

where, 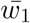 captures the degree-weighted reduction of the adjacency matrix of feeding-links between the primary consumer and basal resources (see appendix 1). In addition, the third term of equation 5 can be reduced to using degree-weighted method as

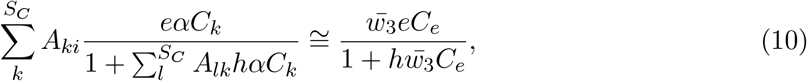

where, 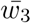 captures the degree-weighted reduction of the adjacency matrix of feeding-links between the secondary consumer and primary consumer (see appendix 1). Finally, the decrease in primary consumer growth rate due to consumption by secondary top predators can be reduced to as (fourth term of equation 4)

This can be done through dimension reduction of the consumers. From the supplementary appendix, after dimension reduction, equation 6 and 3 can be written as:

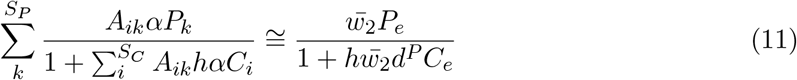

where, 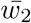 captures the degree-weighted reduction of the adjacency matrix of feeding-links between the secondary consumer and primary consumer (see appendix 1). Thus after degree-reduction high-dimensional equation 4 and 5 can be transformed to

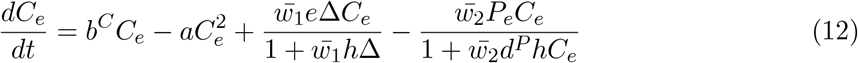

And, the secondary consumer equation is:

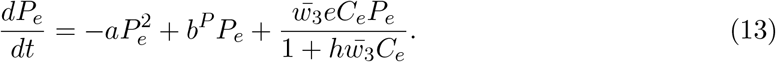

With equation 12 and 13 for a certain perturbation density Δ of the basal producer species, one can evaluate the fixed points of the 2-D equations and thus evaluate whether the entire given food web could be recoverable. In other words, for a given Δ, do the nullclines of equation 12 and 13 intersect at a higher density and thus a stable fixed point is reached? To be noted that, if the two equations intersect at a relatively high density, the recoverability would be larger which we observe in Fig. 2E. For a simulated twelve-species food web in Fig. 2A-D, when Δ = 0.1, i.e., all basal species were controlled to have a density of 0.1, the food web recoverability was not successful (Fig. 2B-C), as the density of consumers and top predators was below 0.01. The 2-D degree-dimension model also correctly characterised this as the nullclines fail to intersect and thus a stable fixed point was not reached (Fig. 2A). When Δ = 5, i.e., all basal species were controlled to have a density of 5, all primary consumers and top predators recover (Fig. 2F, 2G). This was also accurately determined by the reduced 2-D model where the nullclines intersect (Fig. 2E).

**Figure 2:**
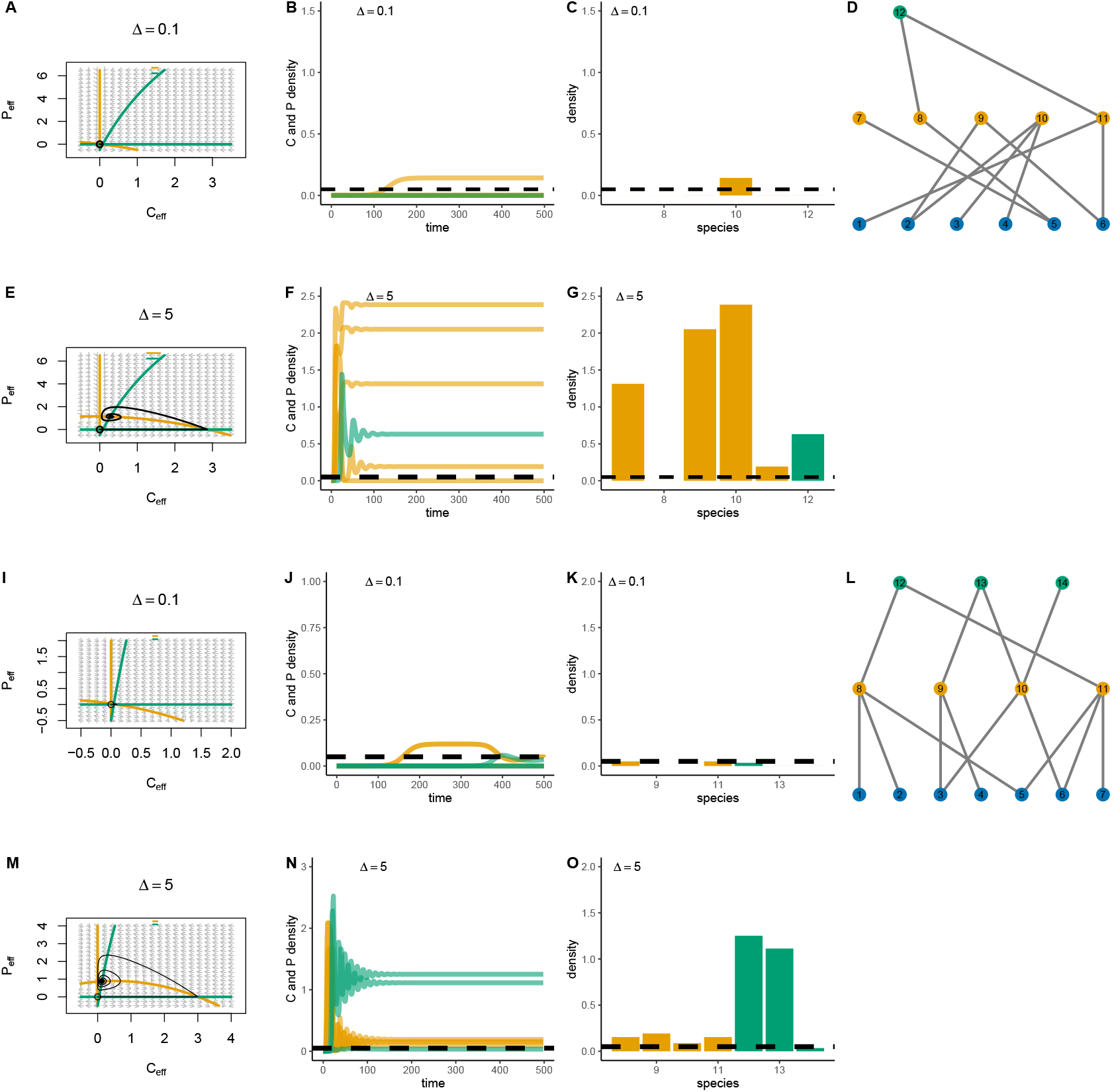
Recovery of collapsed food webs through dimension-reduction and perturbation. (A-D) For a smaller 12 species food web with three trophic levels (D), for a perturbation of Δ = 0.1, the nullclines of the reduced model predicts no appearence of a fixed-point and thus no intersection of the nullclines of *C*_*e*_ and *P*_*e*_. (B-C) Dynamical simulations of equation 4 and 5 show this and equilibrium densities shown by bar plots on C also show this, that no primary or secondary consumer species recover. (E-G) For the same food web, a slightly higher perturbation of the lowest trophic levelΔ = 5 leads to the emergence of a fixed point shown by the intersection of the nullclines and the vector field of the reduced model, which also is observed by the dynamical simulations of equation 4 and 5 shown in F and G. (I-O) For a different tri-trophic food web with 14 species but with three top predators, we also observe similar results. For the dynamical simulations of equation 4 and 5, the parameters are taken from table 1.

Our 2-D reduced model fairly accurately captures the equilibrium dynamics of food webs with varying structure and with temporal white noise (Fig. 3A-F). For a food web with 12 species with one top predator, as Δ increases equilibrium average density of the food web was accurately quantified by the reduced model when compared with average equilibrium density from dynamical simulations of the full model (Fig. 3B-F). Similarly, for another food web with three top predators, the reduced model accurately captures the equilibrium density of full dynamical model (Fig. 3G-L). In comparison to continuous rate of forcing, the 2-D reduced model fairly captures the recovery dynamics of simple 12 species food web (see Fig. S2). In addition, we also tested our 2-D reduced model against two different food webs generated by PMN method. We found that the dimension reduced model again fairly captures the equilibrium densities of the recovery dynamics of the full model of complex food webs with temporal white noise (see Fig. S3-S4).

**Figure 3:**
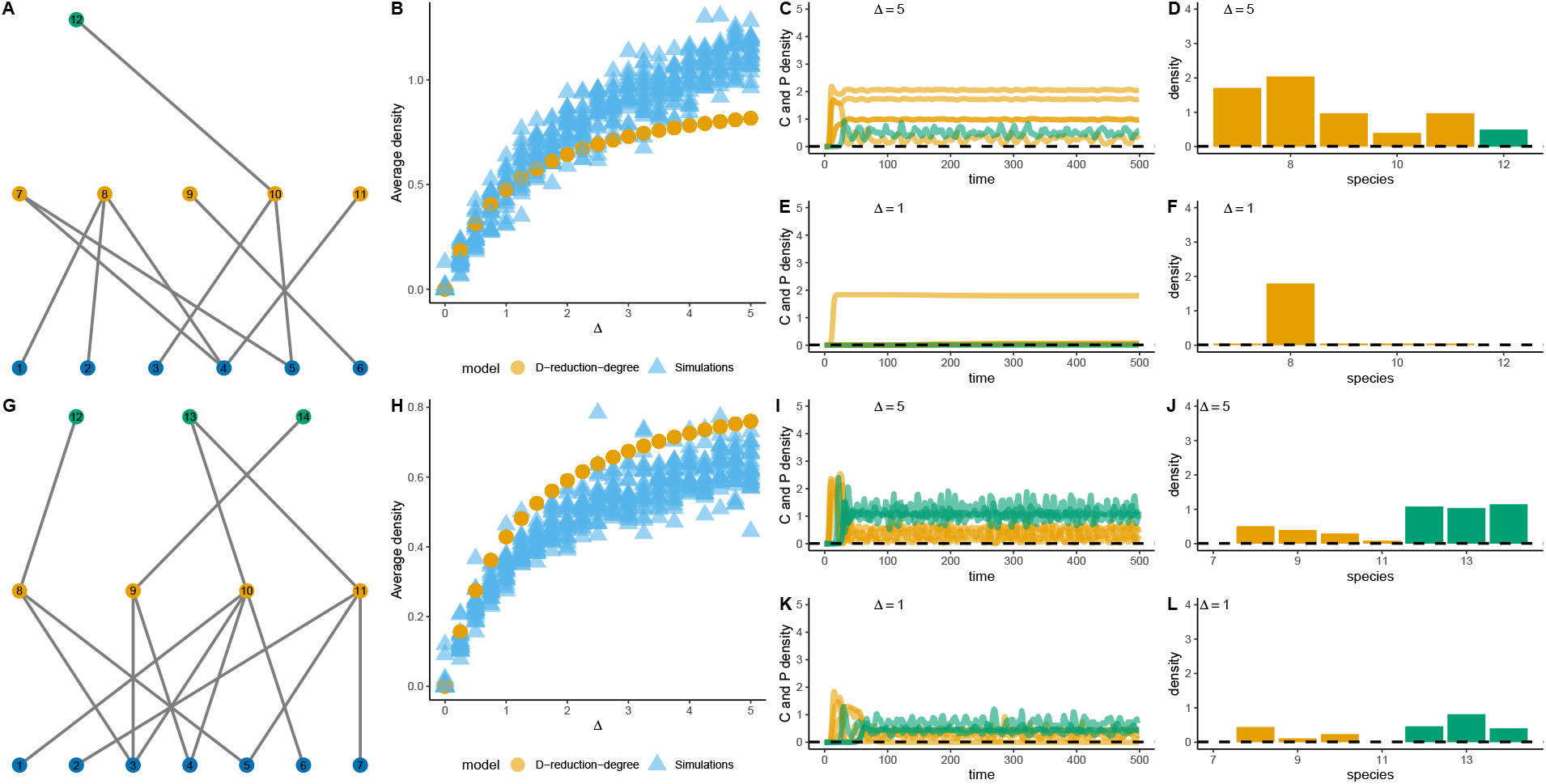
Comparison of recovery dynamics of the dimension-reduced model and dynamical simulations of the full model with temporal white noise, drawn from random normal distribution of mean 0 and variance of 0.1. (A-B) For a food web of 12 species with one top predator, effective equilibrium density of the reduced model (yellow circles) captures the equilibrium density of the full model (blue triangles) very well. Each blue triangle is a replicate simulation of the dynamical model i.e. equation 2 and 3 with temporal white noise.(C-F) For high Δ = 5 all consumer and secondary consumer species recover. For low Δ = 1, only a few species recover. (G-H) For a different tri-trophic food web of 15 species with a connectance of 0.2, effective equilibrium density of the reduced model (yellow dotted circles) for different Δ quite well captures the equilibrium density of the full dynamical food web model (blue triangles). (I-L)Low Δ = 1 leads to only a few species recovery, whereas high Δ = 5 leads to full recovery of the food web. For all the simulations and the reduced model, initial starting density of all primary and secondary consumers were fixed below a density of 0.000001. When there is no perturbation of the lowest trophic level Δ = 0, the system stays at the collapse state. For the parameter values please refer to table 1.

### 3.2 Proportion of predation links, food web connectance on recovery dynamics of food webs

Whether theoretical food webs could be recoverable by perturbing the lowest trophic level were well predicted by the dimension-reduction method. We however did not sample large enough food webs to ensure whether that was universally true. Overall, we found that as the number of predator links increases along with connectance, the dimension-reduced model does poorly in relation to the simulation of the full-model (Fig. 4). This result was however modulated by Δ. As perturbation on the lowest-trophic level Δ increases, the difference in equilibrium density of the reduced model and the full-model increases. In general, at high Δ, high food web connectance and top-predator links, the difference in equilibrium density given by the reduced-model and the full-model simulations increase. This indicates that in such regions of parameter space, the reduced-model might not be a good methodology to assess the recovery dynamics of tri-trophic food webs in general. With PMN food webs we also found qualitatively that if the number of top predation links increases, full recovery of 24 species food web was not achieved (see Fig. S4 E-F).

**Figure 4:**
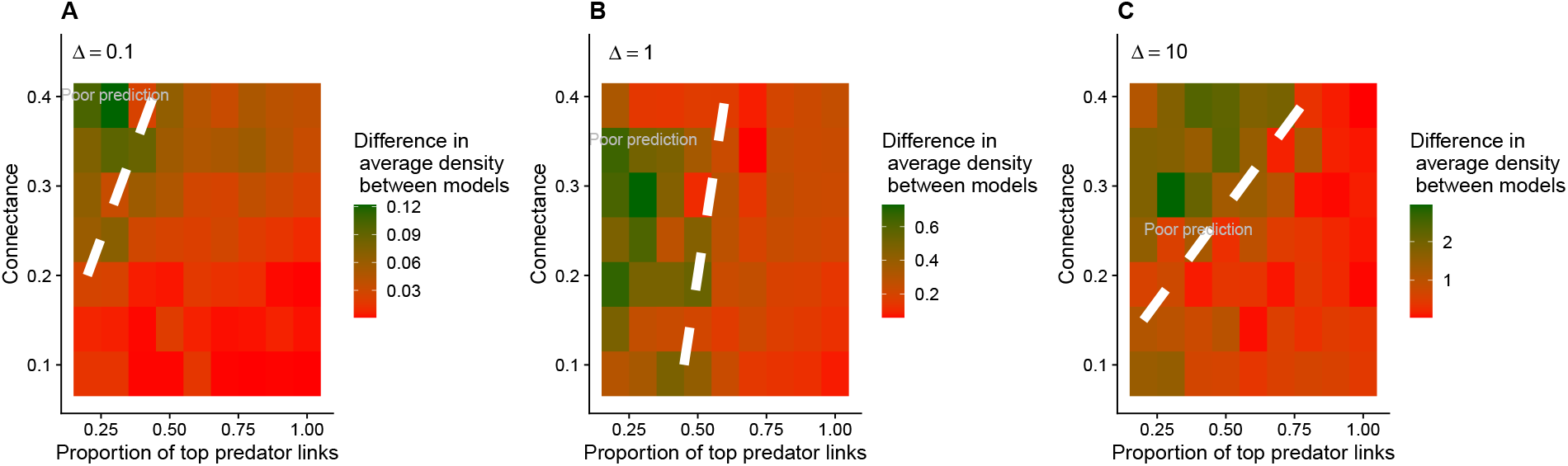
Difference in average equilibrium effective density of the reduced model and full model simulations for different levels of connectance and proportion of top-predator links. For the full model simulations, we took the average equilibrium density across all species. The food web model used was produced through the pyramidal method, with 24 species and the ratio of species in the three trophic levels were of 5:3:2. A) For low Δ = 0.1, there was no big difference in the effective equilibrium density of the reduced model and the full dynamical model. (B-C) But as Δ increases to 1 and 10, the difference in the equilibrium density from the reduced model and the full dynamical model increases. However, at low connectance values, the reduced model still predicts the recovery dynamics of the full dynamical model quite well.The white dashed lines are drawn qualitatively to separate where the effective density of the reduced model diverges from average density from the full model simulations.

### 3.3 Food web connectance, number of basal species forced on recoverability

In general, through simulations of the full model, we show that food webs with high connectance are recoverable from a collapse state if the proportion of top-predator links were low i.e. *<* 0.3 (Fig. 5A-C). This result is also modulated by the amount of perturbation applied to the lowest-trophic level i.e., Δ, with moderate amount of Δ *<* 5 does lead to better recovery of collapsed food webs than high Δ = 5 (Fig. 5A-C).

**Figure 5:**
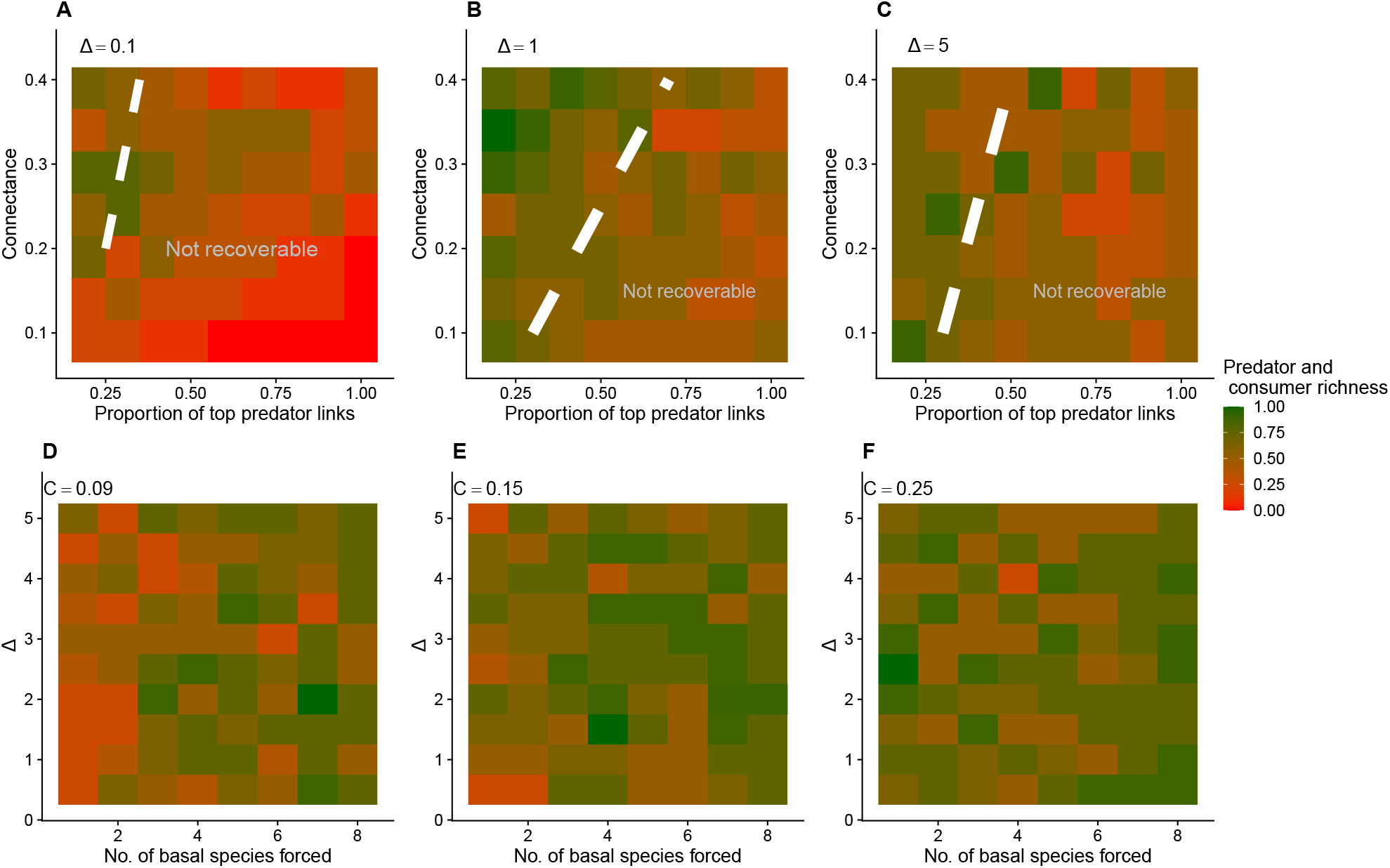
Parameter space and sensitivity analyses of the full dynamical model of tri-trophic food webs of 24 species that differ in connectance and proportion of top predator feeding links for different levels of Δ. (A-C) With larger perturbation Δ = 1 and Δ = 5 food webs with lower connectance and high connectance are recoverable if proporation of predation links are low. (D-F) Number of basal species perturbed also matter in recovery of tri-trophic food webs of different connectance. Specifically, when all 8 basal species are positively perturbed, recovery of tri-trophic food web of 24 species with three difference connectance of 0.09, 0.15, 0.25 was possible. Parameters are from table 1.

In addition, when we evaluated the parameter space of amount of forcing of a food web with 24 species, and number of basal species forced at a time, we found that in general, if all basal species (8 species) of the lowest trophic level was forced, recovery of collapsed the 24 species collapsed food web was possible regardless of the strength Δ (Fig. 5D-F). Furthermore, if the 24 species food web had a higher connectance, recovery was positively impacted but only when number of basal species being perturbed was greater than 6 (Fig 5D-F).

## Discussion

A robust food web can buffer against external perturbations and maintain stability even when changes in a species density could ripple through other trophic levels i.e., trophic cascades (Dunne & Williams 2009, Staniczenko *et al*. 2010). However, as external environmental pressure increases beyond a certain threshold, species extinctions can eventually cascade and lead to collapses at the level of the entire food web (Pinsky *et al*. 2011, Hutchings & Reynolds 2004). Forecasting such extinctions is vital for conservation efforts. While much attention has been given to preventing and forecasting such collapses, less explored is the potential for recovery of collapsed food webs. It remains uncertain whether collapsed food webs can be fully restored and whether some are more recoverable than others (Yacine *et al*. 2021, Dobson *et al*. 2009). Understanding the structure of a food web, characterized by species interactions, could offer insights into the recovery of such collapsed food webs. Here using theoretical food webs, we show that whether complex tri-trophic food webs could be recoverable can be predicted by a dimension-reduced model. Reducing the dimension of complex food webs could greatly help in understanding the overall dynamics, and recovery from a collapse state. We further showed through simulations that amount of perturbation on the lowest trophic level, structure of the food webs, and proportion of predator links impact the recovery of collapsed food webs.

Dimension reduction method have been used to predict tipping points in bipartite networks such as mutualistic networks (Jiang *et al*. 2018, Gao *et al*. 2016). We use similar methodology to reduce the dimensions of a tri-trophic food web and evaluate whether a reduced model of a complex tri-trophic food web can accurately capture the dynamics and thus use that to predict whether recovery from a collapse state is possible. This dimension-reduction method would thus require only minimal amount of species-based information compared to parameterising a whole food web. In addition, a complex multi-species food webs has many different stable and unstable states, reducing the dimension could also further benefit from capturing the stable and unstable states. In our study, the starting point of our modelling framework was the assumption of collapsed food webs i.e., all *P*_*i*_, *C*_*i*_ *<* 0.0001. We then tried to evaluated whether intermediate consumers and top predators of theoretical tri-trophic food webs could recover if the lowest trophic level was restored. We evaluated this through two different avenues: one using a reduced model, and the other through simulations of the entire food web. We thus found that the dimension-reduction model quite accurately captures the recovery dynamics of tri-trophic food webs (Fig. 2,Fig. 3). Perturbing the lowest trophic level at low Δ = 0.1, the reduced-model shows that the nullclines do not intersect, indicating that when the lowest trophic level was kept at Δ = 0.1, there is no fixed point, indicating that recovery was not possible. Indeed, this was similar when the whole food web was simulated (Fig 2A-C). However, if Δ = 5 is increased, i.e., the lowest-trophic level was kept at a higher density, there was an appearance of fixed point as the nullclines of the reduced model intersected, indicating the possibility of recovery, which was simultaneously shown by the simulations of the whole food webs (Fig. 2E-G). Our results thus indicated that the reduced model could be indeed be used to understand the recovery dynamics of a complex food web.

Indeed, the reduced model captured the recovery dynamics of collapsed food webs rather well (Fig. 3). However, at higher values of equilibrium density of the basal species i.e., Δ *>* 1, the difference between the equilibrium effective density of the reduced model and the dynamic simulations increases (Fig. 4C). This indicated an impact of the structure and the amount of perturbation on recovery dynamics of food webs.

Previous studies have demonstrated the bottom-up effects can be modulated by an external environmental pressure that impacts an intermediate consumer of a trophic food web, thereby impacting the net bottom-up impacts in structuring a zooplankton community (Lynam *et al*. 2016). Bottom-up effects of phytoplankton on marine communities have been consistently reported. For instance, Menge *et al*. (1997) showed that bottom-up effects defined by higher growth of phytoplankton could lead to higher food availability for filter feeders and thus further increase denser top predator populations. Thus, in complex marine food webs, recovery by restoring higher densities of basal phytoplankton could lead to recovery of collapsed food webs in general. Our study further adds to this by showing that restoring higher densities of basal resource could not always work and concurrently depends on the structure and proportion of predation links in the community.

The disappearance of apex predators from both aquatic and terrestrial habitats stands out as one of the most significant influences of human activity on ecosystems. Previous research suggests that alterations in the topmost trophic level could trigger top-down cascades and impact ecosystem functions and services provided by complex food webs (Estes *et al*. 2011, Ripple *et al*. 2014). Top-down effects of predation underscore the importance of reinstating or preserving predation dynamics as part of ecological restoration efforts (Ordiz *et al*. 2013, Estes *et al*. 2011). Previous empirical studies have demonstrated that predation could be an important factor in regulation vegetation productivity (Ripple *et al*. 2014, Silliman & Bertness 2002), decreasing the level of eutrophication (Carpenter *et al*. 1999), or maintaining biodiversity in large ecosystems (Beschta & Ripple 2009). Predation offers a top-down control when a complex ecosystem collapses to a state where there is an abundance of the lowest trophic level such as algal blooms in lakes. However, in our study, we considered a bottom-up approach where we assumed that the density of intermediate consumers and top predators are very low such that without any kind of perturbation the whole food web system remains in the undesirable state of collapse.

Our model simulations and the simplified model should be regarded not as definitive truths, but as hypotheses to be empirically tested in small-scale studies. Our models incorporated numerous assumptions, including the absence of omnivory links and the restriction to only three trophic levels. The inclusion of omnivory links could significantly influence the recovery dynamics of theoretical food webs, particularly concerning the recovery of top predators. Moreover, empirical food webs typically encompass more than three trophic levels, which could further complicate extrapolations to the simplified model. With *n* trophic levels, the dimension reduction will lead to atmost *n* − 1 effective equations leading to more stable and unstable states which would be difficult to analytically tract and solve. While it is possible to numerically evaluate the stable and unstable states from three or more equations, it would not be different than just simulating entire food webs. In general, our theoretical study generated three key results: a reduced model could effectively capture complex tri-trophic food web recovery dynamics; lower connectance but a high number of predation links could negatively impact the recovery of food webs; higher number of basal species restoration is key to recovering complex food webs regardless of high or low food web connectance. We argue that recovery of collapsed food webs could be achieved by using a bottom-up approach to restoration, and that simple dimension reduction approach could very well be used to predict the recovery of complex food webs.

## Acknowledgments

GB would like to acknowledge DFG Walter Benjamin grant no BA 7974/1-1 for funding the research. The authors would also like to thank Meike Wittmann for her valuable comments on the manuscript.

## Author contributions

GB formulated the study. SP and GB analysed the model. GB and SP wrote the manuscript.

