## Appendix for "Predicting recoverability of collapsed food webs through perturbation and dimension reduction"

Gaurav Baruah[1], Swastik Patnaik[1], Meike Wittmann[1]

### 1. Degree-based dimension reduction

Using degree-weighted method, the predation of primary consumer on the basal species (third term of equation 4), can be written as

$$\sum_{l}^{S_N} A_{il} \frac{e\alpha\Delta}{1+\sum_{j}^{S_N} hA_{ij}\alpha\Delta} \cong \frac{\bar{w_1} e\Delta}{1+h\bar{w_1}\Delta}, \tag{1}$$

where, $\bar{w_1}$ captures the degree-weighted reduction of the adjacency matrix of feeding-links between the primary consumer and basal resources. We can write how we arrive at $\bar{w_1}$ as follows: first effectively representing each predation on the basal species by the primary consumer as,

$$\sum_{l}^{S_N} A_{il} \frac{e\alpha\Delta}{1+\sum_{j}^{S_N} hA_{ij}\alpha\Delta} = \frac{d_i \alpha e\Delta}{1+d_i h\alpha\Delta}$$

,

Using the degree-weighted method [2], i.e., we weight by the degree of the trophic species in question we arrive at,

$$\bar{w_1} = \frac{\sum_{i}^{S_C} d_i \alpha}{\sum_{i}^{S_C} d_i}. \tag{2}$$

where $d_i$ is the degree of each basal species in the tri-trophic network or number of links each basal species has to the primary consumer, $\alpha$ is the attack rate, $e$ is the conversion efficiency.

In addition, the third term of equation 5 can be reduced to using degree-weighted method as

$$\sum_{k}^{S_C} A_{ki} \frac{e\alpha C_k}{1+\sum_{l}^{S_C} A_{lk} h\alpha C_k} \cong \frac{\bar{w_3} eC_e}{1+h\bar{w_3} C_e}, \tag{3}$$

where, $\bar{w_3}$ captures the degree-weighted reduction of the adjacency matrix of feeding-links between the secondary consumer and primary consumer. We can write how we arrive at $\bar{w_3}$ as follows: first effectively representing each predation on the primary consumer by the secondary consumer or the top predator as

$$\sum_{k}^{S_C} A_{ki} \frac{e\alpha C_k}{1+\sum_{l}^{S_C} A_{lk} h\alpha C_k} = \frac{d_i^C e\alpha C_e}{1+hd_i^C \alpha C_e} \tag{4}$$

where $C_e$ is the effective consumer density, and $d_i^C$ is the number of unique links of the consumer to the top predator. Finally, using degree-weighted method we arrive at,

$$\bar{w_3} = \frac{\sum_{i}^{S_P} d_i^C \alpha}{\sum_{i}^{S_P} d_i^C}. \tag{5}$$

where, the summation goes over all the primary consumer species.

Finally, the decrease in primary consumer growth rate due to consumption by secondary top predators can be reduced to as (fourth term of equation 4). This can be done through dimension reduction of the consumers. From the supplementary appendix, after dimension reduction, equation 4 and 5 can be written as:

$$\sum_{k}^{S_P} \frac{A_{ik}\alpha P_k}{1+\sum_{i}^{S_C} A_{ik} h\alpha C_i} \cong \frac{\bar{w_2} P_e}{1+h\bar{w_2} d^P C_e} \tag{6}$$

where, $\bar{w_2}$ captures the degree-weighted reduction of the adjacency matrix of feeding-links between the secondary consumer and primary consumer. We can write how we arrive at $\bar{w_2}$ as follows: first effectively representing each feeding interaction on the primary consumer by the secondary consumer or the top predator as,

$$\sum_{k}^{S_P} \frac{A_{ik}\alpha P_k}{1+\sum_{i}^{S_C} A_{ik} h\alpha C_i} = \sum_{k}^{S_P} \frac{A_{ik}\alpha P_k}{1+hd_k^P \alpha C_e} = \frac{d_i^C \alpha P_e}{1+hd^P \alpha C_e} \tag{7}$$

where $P_e$ is the effective secondary consumer or top predator density, and $d_i^C$ is the number of unique links of the consumer to the top predator. Finally, using degree-weighted method we arrive at,

$$\bar{w_2} = \frac{\sum_i^{S_C} d_i^C \alpha}{\sum_i^{S_C} d_i^C}. \tag{8}$$

Thus, we then arrive at the effective consumer and top predator equations:

$$\frac{dC_e}{dt} = b^C C_e - aC_e^2 + \frac{\bar{w_1} e \Delta C_e}{1 + \bar{w_1} h \Delta} - \frac{\bar{w_2} P_e C_e}{1 + \bar{w_2} h d^P C_e} \tag{9}$$

And, the secondary consumer equation is:

$$\frac{dP_e}{dt} = -aP_e^2 + b^P P_e + \frac{\bar{w_3} e C_e P_e}{1 + h\bar{w_3} C_e}. \tag{10}$$

**2. Continuous rate of forcing for basal resource species**

Dynamics of tri-trophic food-webs are given by generalised consumer-resource equations [1] with type-2 functional curves as and continuous rate of forcing is :

$$\frac{dN_i}{dt} = N_i \left( b_i^N - \sum_{j=1}^{S_N} a_{ij}^N N_j - \sum_{k=1}^{S_C} \frac{A_{ik} \alpha C_k}{1 + \sum_i^{S_N} h A_{ik} \alpha N_i} \right) + \beta N_i \tag{11}$$

$$\frac{dC_i}{dt} = C_i \left( b_i^C - \sum_{j=1}^{S_C} a_{ij}^C C_j - \sum_{k=1}^{S_P} \frac{A_{ik} \alpha P_k}{1 + \sum_i^{S_C} A_{ik} h \alpha C_i} + \sum_{l=1}^{S_N} \frac{e A_{il} \alpha N_l}{1 + \sum_{j=1}^{S_N} h A_{ij} \alpha N_j} \right), \tag{12}$$

$$\frac{dP_i}{dt} = P_i \left( b_i^P - aP_j + \sum_{k=1}^{S_C} A_{ki} \frac{e \alpha C_k}{1 + \sum_{l=1}^{S_C} A_{lk} h \alpha C_k} \right), \tag{13}$$

With this continuous rate of forcing $\beta$, we evaluated how our 2-D reduced model compared with full model dynamical simulations with continuous forcing. Note that our reduced 2-D model did not incorporate the continuous rate of forcing. This is because, incorporating a continuous rate of forcing would entail solving 3 coupled non-linear equations which is analytically difficult.

Thus our 2-D dimension reduced equations remain same as in the main-text as :

$$\frac{dN_i}{dt} = 0, \tag{14}$$

$$N_i(t) = \beta \tag{15}$$

$$\frac{dC_e}{dt} = b^C C_e - aC_e^2 + \frac{\bar{w_1} e \beta C_e}{1 + \bar{w_1} h \beta} - \frac{\bar{w_2} P_e C_e}{1 + \bar{w_2} h d^P C_e} \tag{16}$$

And, the secondary consumer equation is:

$$\frac{dP_e}{dt} = -aP_e^2 + b^P P_e + \frac{\bar{w_3} e C_e P_e}{1 + h\bar{w_3} C_e}. \tag{17}$$

We thus than vary $\beta$ and then evaluate how full model dynamical simulations i.e., equation 11-13 compares with 2-D reduced model dynamics i.e., equation 16-17. See figure S2.

**3. Probabilistic niche-based food webs (PMN):**

We also created PMN food webs based on the method detailed in Williams *et al.* [5]. This was to test whether our dimension reduction method was able to capture dynamics of food webs generated through PMN method. We thus tested this for first two different food webs one with 16 species and one with 12 species (see Fig. S3,S4). We find that our 2-D dimension reduction method captures the dynamics of food webs generated by PMN webs quite accurately.

**4. Introducing noise in full model dynamics:**

The dynamics of food webs with environmental white noise can be written as following from Ruokolainen & Fowler [4], Ripa & Lundberg [3]:

$$N_i = \Delta \tag{18}$$

$$\frac{dC_i(t)}{dt} = C_i(t)\left(b_i^C - \sum_{j=1}^{S_C} a_{ij}^C C_j(t) - \sum_{k=1}^{S_P} \frac{A_{ik}\alpha P_k(t)}{1+\sum_i^{S_C} A_{ik}h\alpha C_i(t)} + \sum_{l=1}^{S_N} \frac{eA_{il}\alpha N_l(t)}{1+\sum_{j=1}^{S_N} hA_{ij}\alpha N_j(t)} + \epsilon_i^C(t)\right), \tag{19}$$

$$\frac{dP_i(t)}{dt} = P_i(t)\left(b_i^P - aP_j(t) + \sum_{k=1}^{S_C} A_{ki}\frac{e\alpha C_k(t)}{1+\sum_{l=1}^{S_C} A_{lk}h\alpha C_k(t)} + \epsilon_i^P(t)\right), \tag{20}$$

Here, $\epsilon_i^{C,P}(t)$, represents environmental white noise being modelled. At smaller time step, $\epsilon_i^{C,P}(t)$, can be represented by discrete time approximately as $\epsilon_i^{C,P}(T)$, where $T = 0, 1, 2, ....$ Thus, $\epsilon_i^{C,P}(T)$, can be approximated as:

$$\epsilon_i^{(C,P)}(T) = k\epsilon_i^{(C,P)}(T-1) + \sigma\psi_i(T-1) \tag{21}$$

Here, $k = 0$, which means zero auto-correlation between time points, $\sigma$ is the strength of noise which was fixed at 0.1 and $\psi_i(T-1)$ is standard normal components that is independent between species.

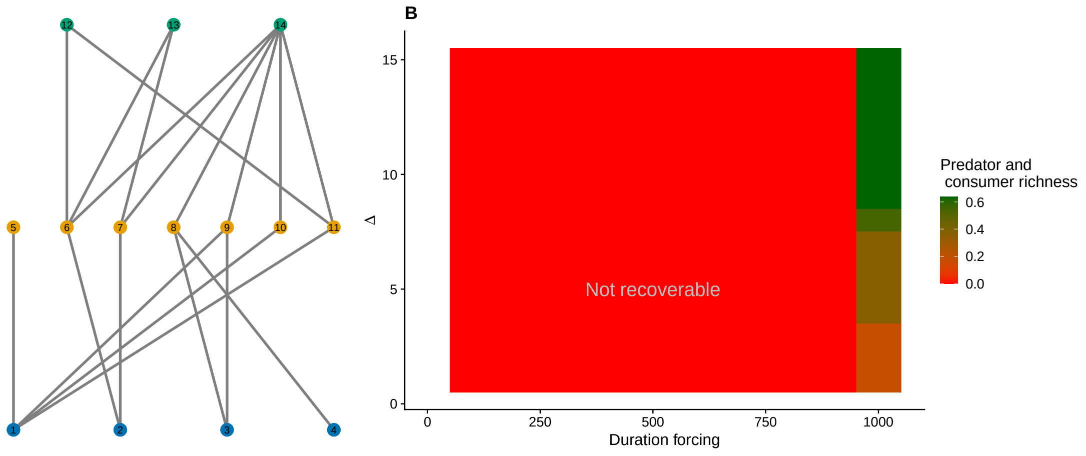


Figure S1: Forcing duration on the basal species and recoverability of a food web with 14 species.

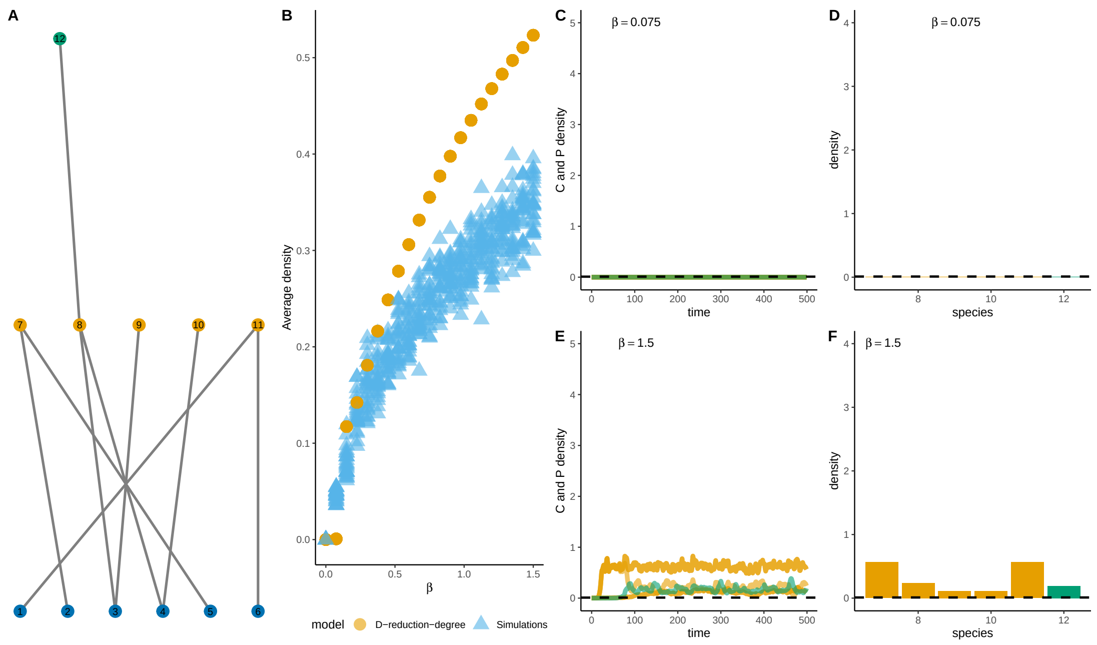


Figure S2: Comparison of recovery dynamics of the dimension-reduced model and dynamical simulations of the full model with temporal white noise and with continuous rate of forcing $\beta$, instead of the $\Delta$. The difference here is that we have a continuous rate of forcing $\beta$, that is linear. We do not fix basal resource density to a constant $\Delta$, instead basal resources can have their own dynamics with an incoming forcing rate of $\beta$. Full model dynamics has temporal white noise drawn from random normal distribution of mean 0 and variance of 0.1. (A-B) For a food-web of 12 species with one top predator, effective equilibrium density of the reduced model (yellow circles) captures the equilibrium density of the full model (blue triangles) very well. Each blue triangle is a replicate simulation of the dynamical model i.e. equation 19 and 20 with temporal white noise. Note that although in the full-model the continuous forcing is given by $\beta$, but the reduced model did not have continuous forcing, but the one in the main-text.(C-F) For high $\Delta = 5$ all consumer and secondary consumer species recover. For low $\Delta = 1$, only a few species recover. (G-H) For a different tri-trophic food-web of 15 species with a connectance of 0.2, effective equilibrium density of the reduced model (yellow dotted circles) for different $\Delta$ quite well captures the equilibrium density of the full dynamical food-web model (blue triangles).

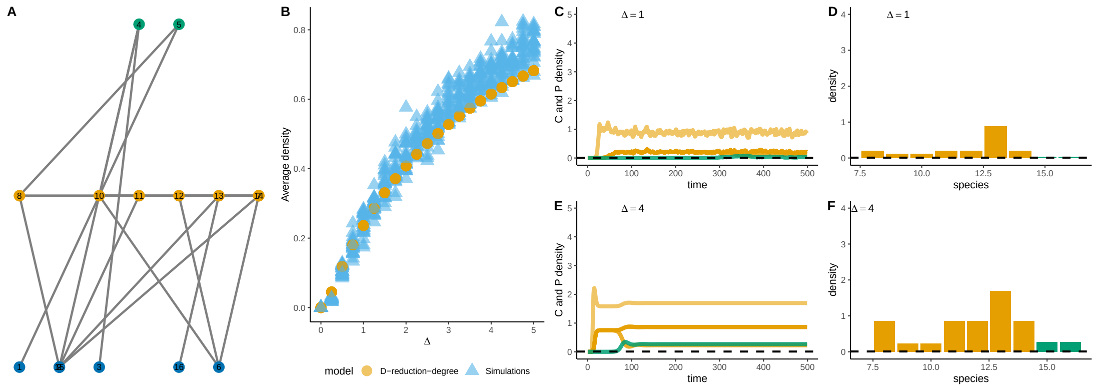


Figure S3: Comparison of recovery dynamics of the dimension-reduced model and dynamical simulations of the full model with temporal white noise for a 16 species food web generated by PMN method. Full model dynamics has temporal white noise drawn from random normal distribution of mean 0 and variance of 0.1. (A-B) For a food-web of 16 species with two predator, effective equilibrium density of the reduced model (yellow circles) captures the equilibrium density of the full model (blue triangles) very well. Each blue triangle is a replicate simulation of the dynamical model i.e. equation 19 and 20 with temporal white noise.(C-F) For high $\Delta = 5$ all consumer and secondary consumer species recover. For low $\Delta = 1$, only primary consumers recover.

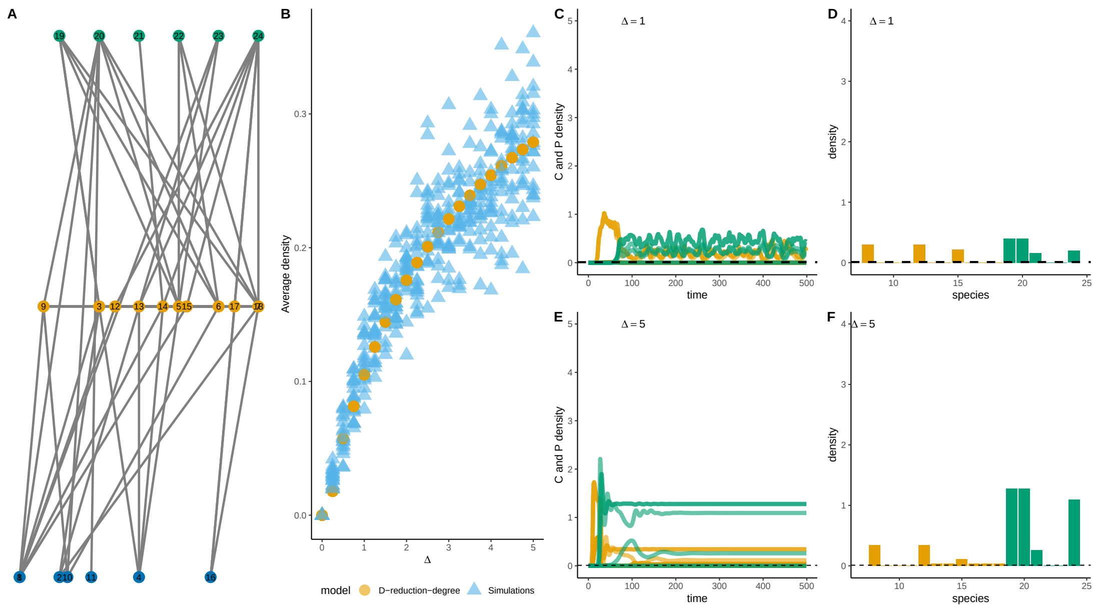


Figure S4: Comparison of recovery dynamics of the dimension-reduced model and dynamical simulations of the full model with temporal white noise for a 24 species food web generated by PMN method. Full model dynamics has temporal white noise drawn from random normal distribution of mean 0 and variance of 0.1. (A-B) For a food-web of 24 species with 6 predators, effective equilibrium density of the reduced model (yellow circles) captures the equilibrium density of the full model (blue triangles) very well. Each blue triangle is a replicate simulation of the dynamical model i.e. equation 19 and 20 with temporal white noise.(C-F) For high $\Delta = 5$ some consumer and secondary consumer species do recover. For low $\Delta = 1$, only few consumer and secondary consumers recover with low density at equilibrium.
